# Exploring The Association of Student Perceptions of Their Teachers’ Science Instruction and Emotional Engagement and The Variance between Grade Level

**DOI:** 10.1101/2024.07.05.602269

**Authors:** Xin Xia, Robert H. Tai

## Abstract

Previous research has consistently shown students have higher learning interests attitudes, and motivation toward science in elementary compared to middle and high school. However, the key predictors behind the observed differences in engagement levels across grade levels remain unclear. This study aims to identify the association between students’ perceptions of science instruction and their emotional engagement, as well as examine the differences across various educational stages. A multilevel random effect model was employed to investigate this association. In addition, the analysis examined whether the associations vary between grade levels. A sample of 6465 students from 25 schools participated in the study.

This study shows that students in higher grades have significantly lower emotional engagement compared to third-grade students. Similarly, students in higher grades have significantly lower values on perceptions of science instruction of interesting science and understandable science decrease compared to third grade. Findings reveal significant associations between students’ perceptions of science instruction, both in perceiving interesting science instruction and understandable science instruction, and their emotional engagement in science learning. The interaction term (Perception of Interesting Science Instruction × Grade) of Grades four, six through eight, and grade 12 are statistically significantly related to emotional engagement, indicating that more positive slopes correspond to high-grade level than third-grade students. The increase in students’ perception of interesting science has the steepest slope of improvement in students’ emotional engagement in 6^th^ grade. We find no statistically significant difference between grades in students’ perception of understandable science and emotional engagement.

Interesting and understandable science classes have positive correlations with students’ emotional engagement in science learning. For future studies, it is worth exploring science instruction during transition grades to better address and mitigate the observed difference in emotional engagement.

## Introduction

Previous research emphasized that students who have a high level of engagement earn positive outcomes in science learning, including academic achievement (Grabau et al., 2017; Lee et al., 2016; Singh et al., 2002), positive attitudes (Osborne et al., 2003; Singh et al., 2002) and interests (Ainley & Ainley, 2011; Renninger et al., 2014), and lifelong learning (Council et al., 2016; Wang & Eccles, 2012). Despite student engagement being an essential component to consider in science education, research has shown that student engagement in science tends to decline as grade levels increase (Jack & Lin, 2017; Wang & Eccles, 2012). Students in higher grades have a lower engagement in science learning.

Wang and Eccles (2012) described that students’ school engagement declined from 7th through 11th grades. They measured three dimensions of student engagement, based on Fredricks et al. (2004) framework, and results showed engagement in all dimensions decreased. Wang and Holcombe (2010) also investigated the relationship between engagement and motivation and attitude of learning. They found that students who disengaged behaviorally and emotionally led to a decrease in their academic motivation, specifically those high-performing students in middle school classes. Additionally, Osborne et al. (2003) also pointed out that the percentage of students pursuing science or science and mathematics declined as well as their attitude and interest in science over 20 years in their review of the literature.

Wang and Eccles (2012) investigated the decline of students’ school engagement using Stage-Environment Fit Theory (Eccles et al., 1993) and suggested that it could be due to fewer opportunities for students to be autonomous and make decisions on their own; less support from teachers; uncaring relationship with teachers; more pressures from comparison when they transition from elementary school to secondary school. Imms and Byers (2017) conducted a study using a single-subject research design in a Queensland school. They discovered that adopting open and flexible classroom layouts, coupled with the integration of one-to-one technology, enhanced students’ perceptions of teaching and learning quality while also boosting student engagement. Similarly, Taylor and Parsons (2011) listed the elements of increasing engagement, including respectful relationships and interaction with teachers; inquiry-based, problem-based, and exploratory activities; relevance to students’ daily lives; multimedia and technology classroom; challenging tasks; and well-designed assessment. However, rare studies questioned students’ perceptions of teachers’ science instruction and how they experience it through grade levels.

Remarkable research has focused on students’ school engagement, however, limited studies investigating the domain of science engagement (Greene, 2015; Schmidt et al., 2018), specifically the potential explanations of engagement decline in science learning. What’s more, relevant fewer studies detect the difference across elementary to high school. To contribute insights into students’ science engagement, this study discovers the association between students’ perception of science instruction specifically on the interesting and understandable science experience from their teachers and students’ science engagement. Meanwhile, this study explores students’ science engagement between different grades (third grade to 12th grade).

### Theoretical Framework

Fredricks et al. (2004) posited three dimensions of engagement: behavioral, emotional, and cognitive. Behavioral engagement involves active participation in learning activities and extracurricular pursuits. Some researchers detected behavioral engagement through students’ attendance and conduct (Christenson et al., 2012; Fredricks et al. 2004), while others measured it through observable behaviors such as reading aloud, raising a hand, discussing with teacher or peers (Ben-Eliyahu et al., 2018). In terms of science learning in K-12 settings, behavioral engagement reflects students’ participation in science activities in formal or informal settings, including but not limited to actively investing time doing science projects, completing science experiments or tasks, etc. Research has indicated a strong correlation between behavioral engagement and achievement in various fields, including science (Schmidt et al., 2018; Sinatra et al., 2015).

Cognitive engagement, characterized by thoughtful involvement and a willingness to invest effort, focuses on understanding complex concepts and mastering difficult skills (Fredricks et al., 2004). It reflects people actively doing something. In the science learning context, cognitive engagement is the willingness to put effort into science learning and participating in science-related activities. Greene (2015) summarized that cognitive engagement in science is a reliable predictor of student achievement in science, building on a review of 20 years of research.

The main focus of this study is emotional engagement. Emotional engagement encompasses feelings of connection to teachers, peers, and school, as well as a genuine interest in learning and enjoyment of problem-solving, linking with students’ attitudes (Fredricks et al., 2004). In science learning content, students enjoy learning science, doing science activities, and interacting with people in formal or informal settings about science. Interests and enjoyment can stimulate and motivate students involved in a science environment as well as formulate students’ identities (Bell et al., 2009). A positive emotion in learning science generally leads students to engage and participate in science activities (Sinatra et al., 2015; Fredricks et al., 2004); and influences their willingness to do work (Connell, J. P., & Wellborn, 1991; Fredricks & McColskey 2012).

### Literature Review Students’ Perception of Interesting Science Instruction

Science is not learned through rote; neither is it accomplished by simply memorizing content or a few formulas. Instead, science learning happens when students are actively engaged in class. An interesting science class is one of the key elements of engagement in learning.

Research has shown that embedding hands-on experiments (Hofstein & Lunetta, 2004), inquiry- based learning (Chen et al., 2014), autonomy and decision-making (Wang & Eccles, 2012), multimedia and technology-enhanced learning environment (Dubovi & Tabak, 2021; Tao et al., 2018) and real-world applications (Bouillion & Gomez, 2001) can captivate students’ interest and engagement in science. For instance, Chen et al. (2014) conducted a longitudinal study to explore the effect of inquiry-based learning on low-achieving students’ attitudes and interests in science. They found that inquiry-based learning can increase students’ attitudes and interest in science supported by classroom observation and interviews. Similarly, Sample McMeeking et al. (2016) investigated students’ perceptions of interest and engagement in science regarding a traveling museum program. They concluded that students’ increased interest in science was due to students’ enjoyment of science and perceived connection to science, rather than a context unrelated to science.

Although research has shown that interesting science classes positively influence students’ attitudes, interests, and engagement in science, it is essential to recognize the significance of students’ perceptions of what makes science interesting. Jack and Lin (2017) also emphasized the significance of comprehending the concept of an interesting science class through the lens of students’ perspectives. Nevertheless, as previous research has shown, interesting science classes contain various factors that are desirable yet sometimes tend to be overlooked in regular science instruction. Students’ perceptions of teachers as well as the enjoyment of science learning contribute to students’ interest in science (Osborne et al., 2003). Some studies explored students’ perceptions of the classroom environment, including teacher support, equity, task orientation (Aldridge et al. 1999; Wolf & Fraser, 2008), different instruction models (Owston et al., 2019), etc., but comparatively less focused on students’ perception of interesting science classes and their perception in different grades.

### Students’ Perception of Understandable Science Instruction

Another key element in engaging students in science instruction is the ability to communicate concepts in a way that students can understand. Clear and concise explanations can facilitate students’ understanding of scientific concepts and further engage in science learning.

Allen and Tanner (2005) highlighted the importance of providing well-structured explanations that avoid technical jargon and incorporate familiar examples to enhance students’ active learning in a large enrollment biology class. Similarly, Osborne et al. (2003) and Aikenhead (2006) emphasized the importance of incorporating real-world applications and addressing contemporary scientific challenges to foster students’ curiosity and sense of purpose in learning science.

Etobro and Fabinu (2017) conducted a study aimed to identify the biological topics in the National Curriculum that Senior Secondary School Two (SSII) students struggle with, understand the reasons behind these difficulties, and propose ways to enhance the effectiveness of teaching and learning in biology. The study collected data from 400 secondary school students. The results highlighted teaching strategies as one reason attributing to the learning difficulties. To address the challenge, the authors suggested using various instructional strategies, and materials; and linking the concepts with students’ daily lives.

Understandable science instruction involves breaking down complex ideas into manageable components and providing real-world examples to illustrate abstract concepts. Previous research has emphasized the importance of teacher practices in facilitating students’ comprehension and engagement in science classrooms. For instance, Hmelo-Silver et al. (2007) emphasized the effectiveness of scaffolding and inquiry-based learning approaches in supporting student achievement, particularly in problem-based learning environments. They advocated that scaffolded inquiry and problem-based settings provide students with opportunities to engage in challenging tasks. However, limited research has investigated the correlation between students’ perceptions of how teachers facilitate understanding in science classes and the extent to which it influences student engagement.

To gain deeper insights into the degree of interesting and understandable science instruction as perceived by students, this study aims to examine students’ perceptions of science instruction across various grade levels and its influence on their emotional engagement. This study can add some nuance in facilitating students’ engagement in science learning.

### Research Questions

1. What is the association between students’ emotional engagement in science and their perceptions regarding teachers presenting science in an interesting way?

a. Does the main effect between students’ perceptions that their teachers present science in an interesting way and their emotional engagement in science vary by grade level?
2. What is the association between students’ emotional engagement in science and their perceptions regarding their teachers presenting science in a clear and understandable way?

a. Does the main effect between students’ perceptions that their teachers present science in a clear and understandable way and their emotional engagement in science vary by grade level?

## Methodology

### Sample

Data used in this study are part of a longitudinal mixed-method study funded by the National Science Foundation (NSF DRL 1811265). These data included 6465 students from grades three to nine within 25 schools. Data in this study were collected at the beginning of the Fall semester of 2012 in public schools located in urban, suburban, town, and rural communities (Author, 2021). In this study, we only use one-year data to identify baseline conditions and provide insights for further investigation in our following longitudinal study.

### Measures

The framework for observing and categorizing instructional strategies (FOCIS) survey is an instrument to measure students’ preferences regarding various types of activities and science engagement (Author, 2021). The survey contains seven dimensions to detect students’ enjoyment of participating in different types of activities. Additionally, the instrument contains self-report questions about science emotional engagement as well as their perceptions about science instruction. Response categories were provided on a 5-point Likert scale, ranging from 1 (Strongly Disagree) to 5 (Strongly Agree). In this study, we focus on the variables of students’ emotional engagement about science learning, and their perceptions of their science instruction.

In terms of demography information, sex is coded 1 for males and 0 for females in these data. The data recorded race and ethnic information into six dummy variables, including White, Hispanic, Black/African American, Asian and Pacific Islander, American Indian, and multi-race. These data contain students from grade three to grade 12. We created dummy variables for each grade.

### Principal Component Analysis (PCA) and Composite Variable

In the FOCIS survey, there are three items to investigate students’ engagement: (1) *Desire*, I have a real desire to learn science; (2) *Favorite*, Science is one of my favorite subjects; (3) *Enjoy*, Science is something I enjoy very much. To create a composite variable for students’ engagement, we conduct a component factor analysis. To effectively combine these three variables into a single measure of student emotional engagement, we employ Principal Component Analysis (PCA), which is a statistical technique used to test the dimensionality of datasets (Jolliffe, 2002).

In the context of our survey, PCA helps in identifying the underlying factor, students’ emotional engagement in science, that explains the maximum variance shared among the three items. The first principal component derived from this analysis is then used to create a composite variable. This composite score represents a weighted sum of the original items, where the weights are the factor loadings of the first principal component (Abdi & Williams, 2010). Using this composite variable, we can more accurately and succinctly quantify and analyze variations in students’ emotional engagement.

### Statistical Analysis

This study runs through the RStudio *lme4* package. We obtained descriptive data for each survey item of a compositive outcome variable by summarizing the variables, which included calculating the mean, and skewness scores, and conducting a Kurtosis test.

### Construct Validity and Reliability

The Cronbach’s alpha of the three items is 0.881 indicating a high level of internal consistency reliability for the three variables (Desire, Favorite, and Enjoy). It suggests that they measure a similar underlying construct of students’ science engagement.

The loadings of the items on each of the three principal components are listed in Table 1, indicating how each item contributes to the principal component. A composite variable is constructed by combining three individual variables: “Desire,” “Favorite,” and “Enjoy.” Each variable is weighted based on its contribution to overall student emotional engagement, with "Desire" weighted at 0.573, "Favorite" at 0.585, and "Enjoy" at 0.574. We maintained the original 5-point Likert scale values using the equation below:

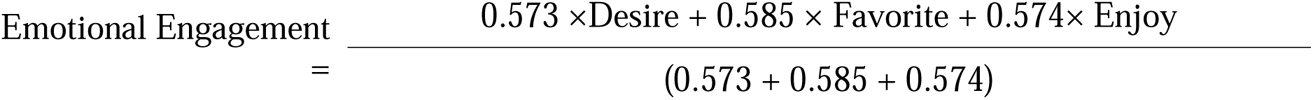

**Table 1:** The principal components analysis (PCA) and descriptive results on emotional engagement.

| Items | Loading | Mean | skew | kurtosis | sd | □ |
| --- | --- | --- | --- | --- | --- | --- |
| Desire: I have a real desire to learn science | 0.573 | 3.24 | -0.23 | -1.27 | 1.45 | 0.881 |
| Favorite: Science is one of my favorite subjects | 0.585 | 3.30 | -0.32 | -1.35 | 1.53 |  |
| Enjoy: Science is something I enjoy very much | 0.574 | 3.42 | -0.40 | -1.11 | 1.41 |  |

*Notes: SE: Standard Error □: Cronbach's Alpha.*

#### Multilevel Modeling

To investigate the relationship between the students’ perceptions of teaching science (clear and understandable presentation and making science interesting) and students’ science engagement, this study employs a random effects model.

In the model, students are nested within schools, which allows us to estimate the variance in science engagement explained by the quality of science teaching (i.e., clear and understandable; interesting) while accounting for potential clustering effects at the school levels. Since students in this study are nested within schools, we use random effect modeling to eliminate the assumption of independence and misestimation of standard errors associated with ordinary least squares regression (Bickel, 2007; Raudenbush & Bryk, 2002).

Model one was the unconditional model and model two was the main effect model which included all independent variables. In model three, we included interaction terms for grade level and students’ perception of teachers making science interesting. Similarly, model four contained the interaction effect of grade and students’ perception of clear and understandable science instruction. To test the model fit, we use a likelihood ratio test (LRT), using a chi-square distribution with the degrees of freedom.

In this study, we used multilevel modeling (MLM). Two variables are used to present the students’ perception of science instruction: (1) *Interesting Science*, My teacher presents science in a clear and understandable way, and (2) *Understandable Science*, My teacher makes science interesting.

The model 2 was defined as follows:

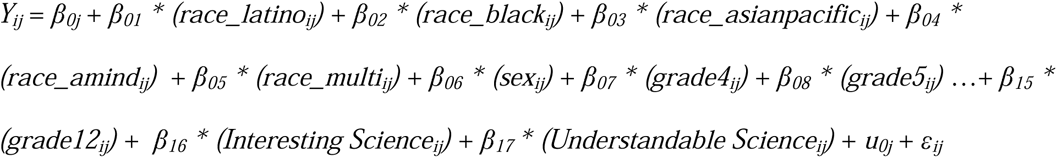

The model 3 (interaction models) was defined as follows:

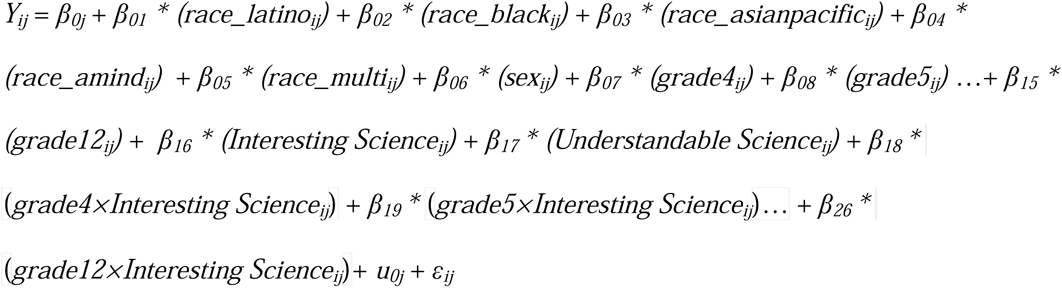

Where:

*Y_ij_* represents the science emotional engagement score for the *i^th^* student in the *j^th^*classroom.

β*_0j_* represents the intercept.

Β*_01-05_*represents the racial and ethnic groups.

Β*_06_* represents the fixed effect of gender.

Β*_07-15_*represents the fixed effect of grades.

Β*_16_* represents the fixed effect of students’ perception of interesting science teaching.

Β*_17_* represents the fixed effect of students’ perception of the clarity of science teaching.

β*_18_* -*_26_* represents the interaction effects of grade and students’ perception of interesting science teaching

*u_0j_* represents the random effect for schools.

ε*_ij_* represents the errors

## Results

### Descriptive Statistics

The data contained roughly equal percentages of participants identifying as male (50.53%) and female (49.47%) from a total of 6465 students. Participants self-reported their racial/ethnic identity in Fall 2012 as: 49.06% White, 17.26% Hispanic, 16.52% African American, 15.19% with multiple racial/ethnic identities, and 1.96% Asian and Pacific Islander. There are similar demographics between 3 and 8^th^-grade participants and grades 9 to 12 contained relatively smaller sample sizes.

### The Descriptive Analysis of Emotional Engagement and Students’ Perceptions of Science Instruction

Table 2 presents the mean scores of student science emotional engagement and the perceived quality of teaching science across different grade levels. In terms of student science engagement, the mean scores range from 2.89 in grade 10 to 3.93 in grade three, showing students in higher grades have lower emotional engagement compared to grade 3. Before sixth grade, students’ scores on emotional engagement are above the grand mean (*M _emotional engagement_* =3.34), while scores drop below the grand mean for students entering sixth grade and above.

**Table 2.** Students’ emotional engagement and the perceptions of science instruction by grade level.

| Grade | Mean of Emotional Engagement | Mean of Teachers making Science Interest | Mean of Teachers Presenting Science Clear |
| --- | --- | --- | --- |
| 3 | 3.93 | 4.35 | 4.34 |
| 4 | 3.69 | 4.27 | 4.24 |
| 5 | 3.74 | 4.29 | 4.27 |
| 6 | 3.24 | 3.84 | 3.86 |
| 7 | 3.21 | 3.88 | 3.79 |
| 8 | 3.10 | 3.76 | 3.82 |
| 9 | 2.97 | 3.26 | 3.30 |
| 10 | 2.89 | 3.37 | 3.38 |
| 11 | 3.06 | 3.56 | 3.65 |
| 12 | 2.95 | 3.39 | 3.41 |

This suggests that students in elementary have relatively higher emotional engagement scores, and students in higher grades have lower emotional engagement scores in science.

Regarding the perception of science teaching, two aspects are assessed: making science interesting and presenting science clearly. As Table 2 shows, students in lower grades generally rate higher scores in perceptions of their teachers making science interesting compared to higher-grade students. This indicates that students perceive more positively in terms of their teachers’ ability to make science interesting in lower grades. However, students in sixth grade or higher level, the means of each grade fall below the grand mean (*M _interesting science_* =3.85). Likewise, students perceive teachers teaching science more clearly in lower grades compared to higher grades in science (Table 2).

### Model Comparing

We conduct a likelihood ratio test (LRT) to examine the model fit with the additional parameters of each new model using the -2-log likelihood (-2LL) statistics. The LRT test uses a chi-square distribution with the degrees of freedom reflecting the number of parameters added to the model. If a model provided a better fit than the previous model, then the null hypothesis was rejected. Compared to model two the chi-square of model three and model two is 28.396 (χ_obe_ ^2^ = 28.396, df=9, p<0.001), which means model three fits better than model two. However, none of the interactions of model four is significant (χ_obe_ ^2^ = 15.685, df=9, p=0.07), which means there is no statistically significant difference in the effect of perceptions of understandable science teaching on emotional engagement between grade levels. Therefore, model 3 is our best model for research question one, and model 2 is the best model for research question two.

### Model Building

Model one is an unconditional model in that no predictors were estimated. Intra-class correlation (ICC) is 0.098 indicating that 9.8% of the variance in students’ science emotional engagement is between schools. Model 1 shows that there was a variation in students’ science emotional engagement scores by schools (τ_00_ = 0.17). The average science emotional engagement score is predicted to be 3.54, statistically different from zero (p< 0.001). Model two contains all predictor variables, including students’ perceptions about science instruction (i. e. interesting science teaching and understandable science teaching), student gender, racial and ethnic group, and grade level.

Model 2 answers research question 2. There is a statistically significant positive association between students’ perception of understandable science teaching and emotional engagement. For each unit increase in students’ perception of understandable science instruction, the emotional engagement score statistically significantly increases by 0.18, holding all other variables constant (*b* =0.18, *CI*= [0.16, 0.21], *p*-value <0.001). Since Model 2 is a better fit than Model 4, meaning the interaction term (perception of understandable science instruction × grade) is not statistically significantly related to emotional engagement. Therefore, to answer sub-question of research two, the main effect is not very by grade level.

Model 3 in Table 3 reveals a statistically significant positive association between students’ perception of science instruction and emotional engagement. To address research question one, Model 3 shows there is a positive association between students’ perception of interesting science teaching and emotional engagement. For each unit increase in perception of interesting science teaching, there’s an associated 0.32 increase in the emotional engagement score statistically significantly, holding all other variables constant (*b*=0.32, *CI*= [0.25, 0.38], *p*- value<0.001).

**Table 3.**
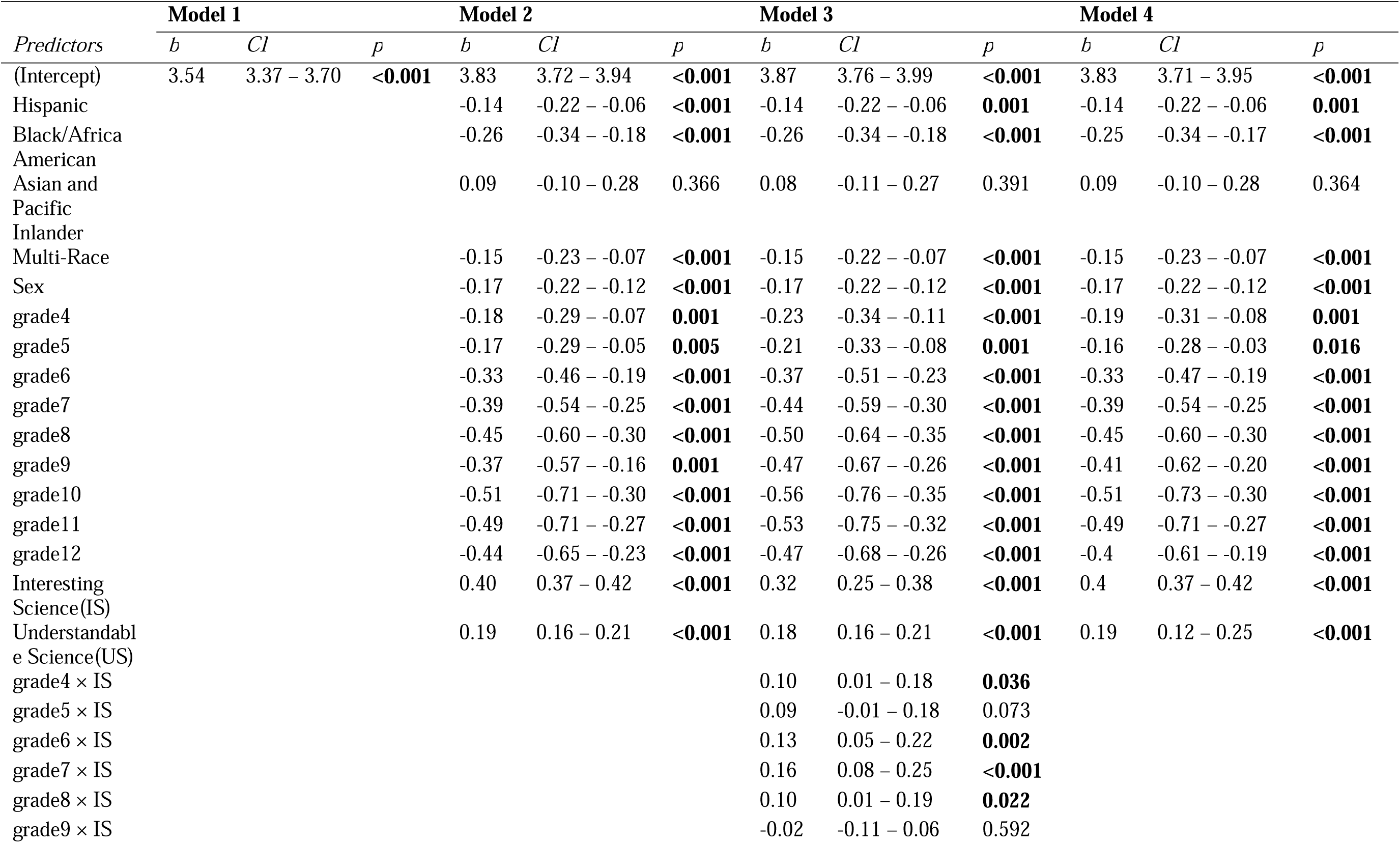
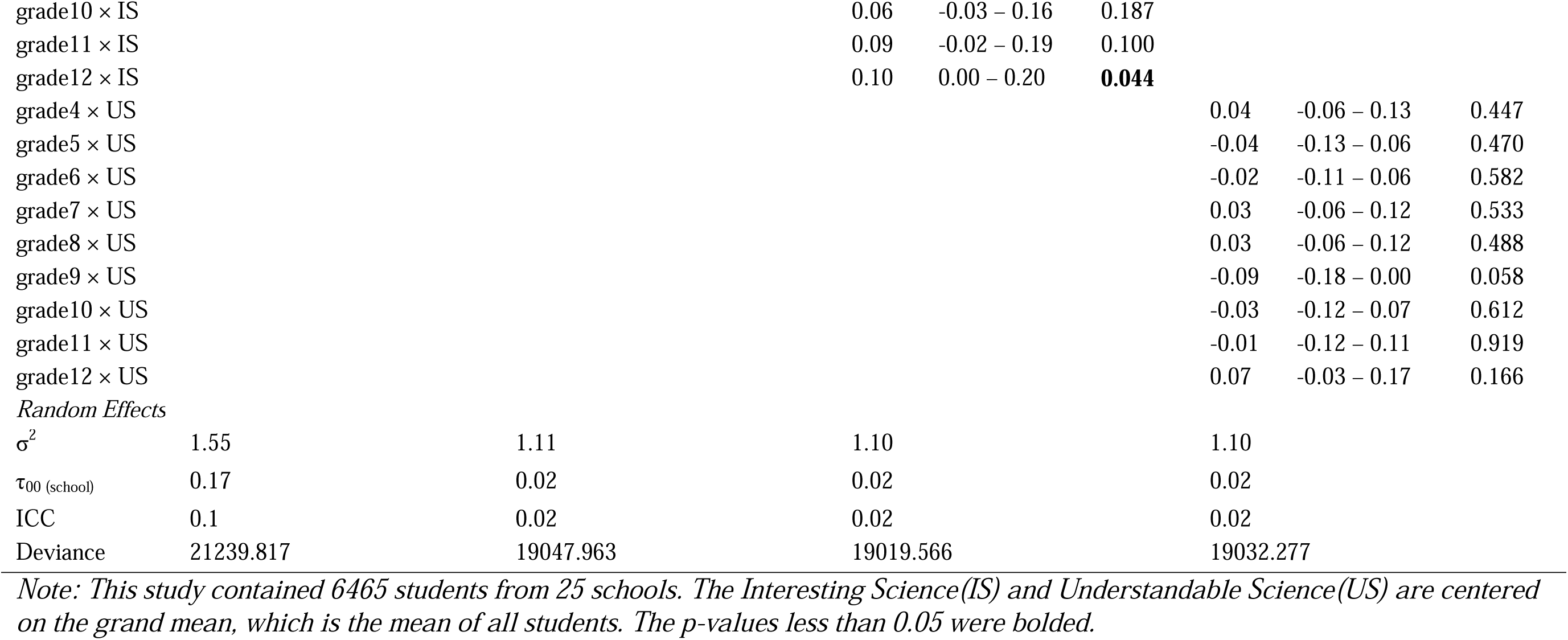
Regression Results.

Model 3 addresses the sub-questions of research question one. The estimated interactions suggest that the relationship between student’s emotional engagement and their perception of interesting science varies in different grade levels. There was a statistically significant interaction between gender 4 and students’ perceptions of interesting science teaching (*b* = 0.095, p<0.05).

Similarly, students in grade 6 (*b* = 0.134, p<0.01), grade 7 (*b*= 0.163, p<0.001), grade 8 (*b*= 0.101, p<0.05), and grade 12 (*b* = 0.100, p<0.05) with higher perceptions of interesting science teaching rate higher emotional engagement but the rate of change is different than third-grade students.

As Figure 1 shows, Grade 6 shows the steepest increase in emotional engagement relative to how interested they feel about the science instruction. This suggests that in the transition year in which students move from elementary to middle school, students’ emotional engagement is most sensitive to their perceptions of how interesting their teachers present the science class. For fourth graders, students’ perceptions of what extent teachers teaching science in an interesting way had a positive association with students’ emotional engagement. However, the rate of increase is relatively smaller than third grade.

**Figure 1.**
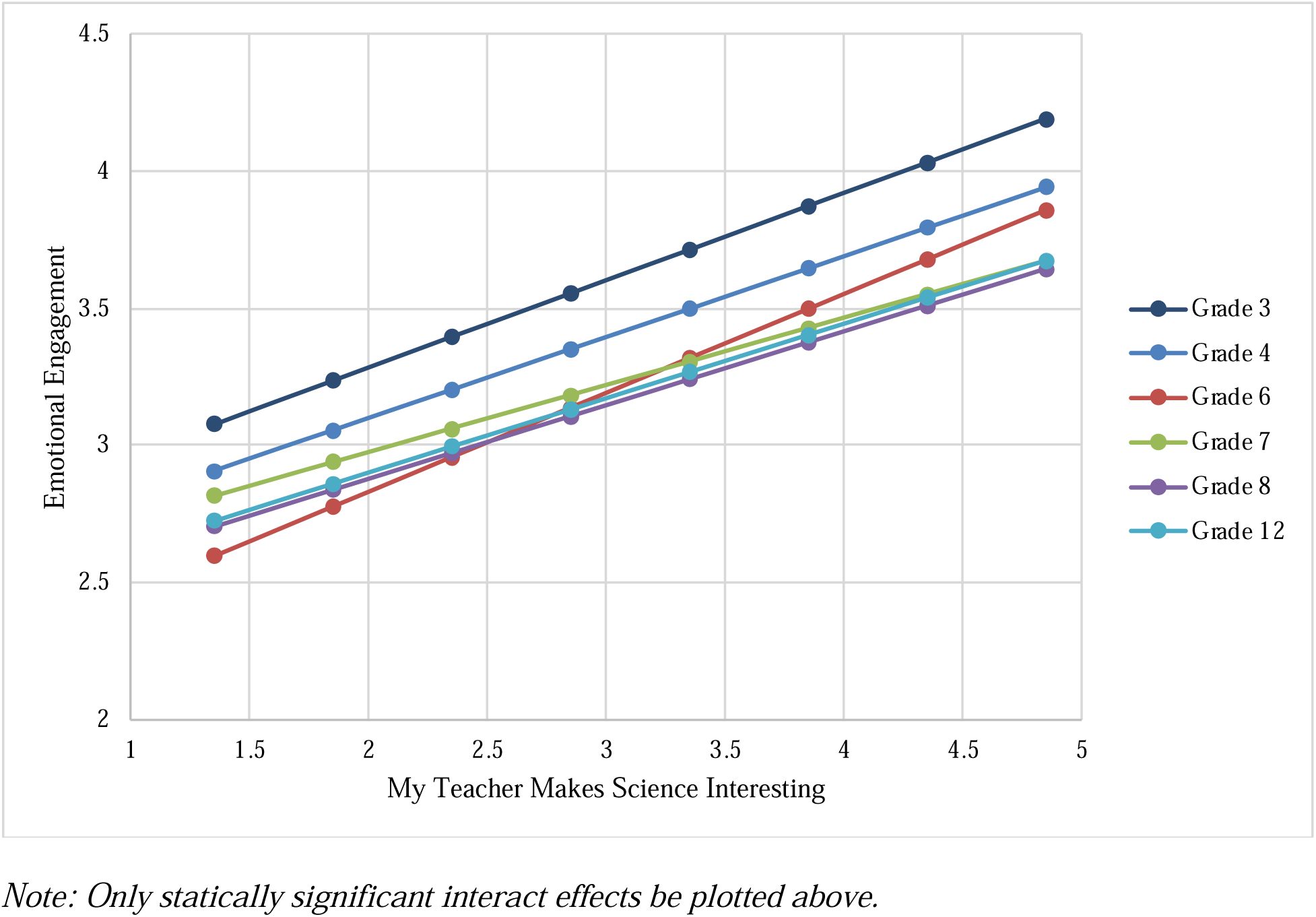
Interaction Effect of Grades and Students’ Perception of Interesting Science Instruction

Grades 7, grade 8, and grade 12 also show a clear positive trend but with a slightly flatter slope compared to Grades 3. This indicates that while making science interesting is important, its direct impact on emotional engagement might be slightly less pronounced than at the transition stages of entering middle school.

## Discussion

The findings of this study shed light on various factors influencing students’ emotional engagement in science learning within the school context. The results underscore that students’ emotional engagement declines while they progress to higher grade levels. The finding aligns with previous research (Jack & Lin, 2017; Osborne et al., 2003; Wang & Eccles, 2012). A sufficient number of studies showed a decline in students’ interest, attitude, and motivation towards learning science as they progress through their academic journey.

Studies have shown that students’ interest and attitude toward science decline from age 11 onwards (Osborne et al., 2003; Potvin & Hasni, 2014; Tröbst et al., 2016; Vedder Weiss & Fortus, 2012). For instance, Potvin and Hasni (2014) emphasized that students’ interests and attitudes toward science tend to fade from the beginning of their education to the secondary level. Furthermore, this study reveals that the decline in students’ emotional engagement begins at an even earlier stage in elementary education. There is a significant gap between the fifth and sixth grades. Prior to sixth grade, students’ scores for emotional engagement are above average, whereas for students moving to sixth grade and above, scores drop below the average of all students scores.

Additionally, students in higher grade levels have lower scores on emotional engagement in science learning compared to grade 3 students. What’s more, the interaction analysis reports that students’ emotional engagement in the transition year (grade 6) is more sensitive about their perception of interesting science classes. Their emotional engagement has a higher improvement if their perception of science class is more interesting compared to third-grade students.

However, to what extent their perception of teachers making science clear and understandable do not find statistically significant differences between grades. This finding emphasizes the importance of addressing potential declines in affective engagement as students progress through the educational system, which can be done through targeted interventions and instructional strategies to maintain students’ interest and motivation to learn science at different grade levels. Meanwhile, it implies that there needs more research on science instruction in the transition grades (Karalar et al, 2021).

Regarding students’ perceptions of science instruction, there is an observed difference in their feelings about receiving interesting science classes and understandable science classes between grade levels. Two gaps in students’ perceptions of science instruction are noticeable: one occurring between fifth and sixth grades and another between eighth and ninth grades. This might be due to the transition from elementary to middle school, which can be a significant change in teaching styles (Ebenezer & Zoller, 1993; Osborne et al., 2003), and assessment pressure (Erdem, 2012; Potvin & Hasni, 2014), etc. Science content has a tendency to become more difficult when students enter middle or high school. Science teachers may be less engaged in designing attractive and interesting science classes. As science content becomes more complex, teachers may face more challenges in breaking down concepts and guiding students to understanding.

Regression modeling reveals that students are more likely to be emotionally engaged in science learning when they have positive feelings about receiving interesting science classes and clear and understandable science classes. This emphasizes the importance of advocating for teachers to prepare science classes in an interesting, organized, easy-to-understand way to foster positive emotional experiences in science classrooms and enhance students’ overall learning outcomes and attitudes toward science. It reflects the importance of teachers’ pedagogical training and practice in science education (Jungert et al, 2020).

## Conclusions

Science instruction plays a key role in students’ engagement in science learning. This study reveals that students have a higher emotional engagement in early elementary. Comparing students from grade three through grade 12, students in higher grades have a lower emotional engagement in science learning in general with a notable gap between fifth and sixth grades.

Prior to this transition, students’ emotional engagement scores were above average, but the scores dropped below average for students in sixth grade and beyond.

The regression modeling underscores that positive perceptions of interesting and clear science instruction are key predictors of emotional engagement, emphasizing the critical role of teachers in designing engaging and understandable science lessons. Additionally, higher grade levels exhibit lower emotional engagement compared to grade 3 students. Interaction analysis indicates that students’ emotional engagement in grade 6 is particularly sensitive to their perceptions of the interesting nature of science classes, suggesting that enhancing the perceived interest in science instruction could mitigate engagement declines during this critical transition period. However, students’ perception of clear and understandable science instruction does not have a significant difference between grade levels.

Overall, the findings of this study contribute to our understanding of students’ emotional engagement and their perception of science instruction between different levels. By identifying predictors of emotional engagement, educators and policymakers can implement targeted interventions and instructional strategies to create interesting learning environments, promote positive affective experiences, and increase overall student engagement and success in science learning. Further research focused on science instruction during transition grades is recommended to better address and mitigate the observed difference in emotional engagement.

## Limitation

One limitation of the study is that the study did not delve into the specific reasons behind students’ perceptions of interesting science classes and clear, understandable science instruction. Understanding these underlying reasons could provide valuable insights into the factors contributing to students’ engagement with science education. Additionally, this study did not explore students’ cognitive and behavioral engagement in science learning. For future research, it is valuable to conduct qualitative interviews or quantitative research about how the students picture interesting science instruction or understandable science in different grade levels.

This study explores the difference in students’ emotional engagement and perception of science instructions between different grades. In the future, longitudinal studies are needed to investigate the trend of students’ perception of science instruction as well as multilevel engagement in science learning.

It is recommended to conduct qualitative interviews or surveys to delve deeper into teachers’ experiences as well. This approach can help uncover the underlying reasons behind their perspectives as science instructors and shed light on potential challenges encountered in science teaching. Furthermore, educators could also benefit from incorporating diverse teaching strategies and approaches based on students’ varied interests and learning preferences, thereby fostering greater engagement and participation in science learning.

## Authors’ contributions

All authors contributed to the study’s conception and design. Material preparation, data collection were performed by Dr. Robert H. Tai, and analysis by Xin Xia. The first draft of the manuscript was written by Xin Xia, and all authors commented on previous versions of the manuscript. All authors read and approved the final manuscript.

## Acknowledgments

Parts of this work have been supported by the US National Science Foundation (NSF DRL 1811265). Any opinions, findings, and conclusions or

recommendations expressed in this material are those of the author(s) and do not necessarily reflect the views of the US National Science Foundation.

## Availability of data and materials

The data used in this research is derived from a project founded by the National Science Foundation under Grant No. (NSF DRL 1811265). The datasets analyzed during the current study are available from the corresponding author upon reasonable request.

## Reference

1. Abdi, H., & Williams, L. J. (2010). Principal component analysis. Wiley Interdisciplinary Reviews: Computational Statistics, 2(4), 433–459.

2. Aikenhead, G. S. (2006). Science education for everyday life: Evidence-based practice.

3. Ainley, M., & Ainley, J. (2011). Student engagement with science in early adolescence: The contribution of enjoyment to students’ continuing interest in learning about science. Contemporary Educational Psychology, 36(1), 4–12.

4. Aldridge, J. M., Fraser, B. J., & Huang, I. T. C. (1999). Investigating classroom environments in Taiwan and Australia with multiple research methods. Journal of Educational Research, 93, 48–57.

5. Allen, D., & Tanner, K. (2005). Infusing active learning into the large-enrollment biology class: seven strategies, from the simple to complex. Cell Biology Education, 4(4), 262–268.

6. Allen, D., & Tanner, K. (2021)

7. Bell , P. , Lewenstein , B. , Shouse , A. and Feder , M. (2009) . Learning science in informal environments: People, places, and pursuits.

8. Ben-Eliyahu, A., Moore, D., Dorph, R., & Schunn, C. D. (2018). Investigating the multidimensionality of engagement: Affective, behavioral, and cognitive engagement across science activities and contexts. Contemporary Educational Psychology, 53, 87–105.

9. Bickel, R. (2007). Multilevel analysis for applied research: It’s just regression!. Guilford Press.

10. Bouillion, L. M., & Gomez, L. M. (2001). Connecting school and community with science learning: Real world problems and school–community partnerships as contextual

11. scaffolds. Journal of Research in Science Teaching: The Official Journal of the National Association for Research in Science Teaching, 38(8), 878–898.

12. Chen, H. T., Wang, H. H., Lin, H. S., P. Lawrenz, F., & Hong, Z.R. (2014). A longitudinal study of an after-school, inquiry-based science intervention on low-achieving children’s affective perceptions of learning science. International Journal of Science Education, 36(13), 2133–2156.

13. Christenson S L., Reschly, A. L., & Wylie, C. (2012). Handbook of Research on Student Engagement.

14. Connell, J. P., & Wellborn, J. G. (1991). Competence, autonomy, and relatedness: A motivational analysis of self-system processes.

15. Council, T. A., & National Academies of Sciences, Engineering, and Medicine. (2016). Science teachers’ learning: Enhancing opportunities, creating supportive contexts. National Academies Press.

16. Dubovi, I., & Tabak, I. (2021). Interactions between emotional and cognitive engagement with science on YouTube. Public Understanding of Science, 30(6), 759–776.

17. Ebenezer, J. V., & Zoller, U. (1993). Grade 10 students’ perceptions of and attitudes toward science teaching and school science. Journal of Research in Science Teaching, 30(2), 175–186.

18. Eccles, J. S., Midgley, C., Wigfield, A., Buchanan, C. M., Reuman, D., Flanagan, C., et al. (1993). Development during adolescence: The impact of Stage-Environment Fit on young adolescents’ experiences in schools and in families. American Psychologist, 48, 90–101.

19. Erdem, E. (2012). Examination of the effects of project based learning approach on students’ attitudes towards chemistry and test anxiety. World Applied Sciences Journal, 17(6), 764–769.

20. Etobro, A. B., & Fabinu, O. E. (2017). Students’ perceptions of difficult concepts in Biology in senior secondary schools in Lagos State. Global Journal of Educational Research, 16(2), 139–147.

21. Fredricks, J. A., Blumenfeld, P. C., & Paris, A. H. (2004). School engagement: Potential of the concept, state of the evidence. Review of educational research, 74(1), 59–109.

22. Fredricks, J. A., & McColskey, W. (2012). The measurement of student engagement: A comparative analysis of various methods and student self-report instruments. In Handbook of research on student engagement (pp. 763–782).

23. Greene, B. A. (2015). Measuring cognitive engagement with self-report scales: Reflections from over 20 years of research. Educational Psychologist, 50(1), 14–30. 10.1080/00461520.2014.989230

24. Grabau, L. J., & Ma, X. (2017). Science engagement and science achievement in the context of science instruction: A multilevel analysis of US students and schools. International Journal of Science Education, 39(8), 1045–1068.

25. Hmelo-Silver, C. E., Duncan, R. G., & Chinn, C. A. (2007). Scaffolding and achievement in problem-based and inquiry learning: a response to Kirschner, Sweller, and. Educational Psychologist, 42(2), 99–107.

26. Hofstein, A., & Lunetta, V. N. (2004). The laboratory in science education: Foundations for the twenty first century. Science Education, 88(1), 28–54.

27. Imms, W., & Byers, T. (2017). Impact of classroom design on teacher pedagogy and student engagement and performance in mathematics. Learning Environments Research, 20, 139–152.

28. Jungert, T., Levine, S., & Koestner, R. (2020). Examining how parent and teacher enthusiasm influences motivation and achievement in STEM. The Journal of Educational Research, 113(4), 275–282.

29. Karalar, H., Sidekli, S., & Yıldırım, B. (2021). STEM in transition from primary school to middle school: primary school students’ attitudes. International Electronic Journal of Elementary Education,13(5)

30. Lee, C. S., Hayes, K. N., Seitz, J., DiStefano, R., & O’Connor, D. (2016). Understanding motivational structures that differentially predict engagement and achievement in middle school science. International Journal of Science Education, 38(2), 192–215.

31. Lin, H. S., Hong, Z. R., & Huang, T. C. (2012). The role of emotional factors in building public scientific literacy and engagement with science. International Journal of Science Education, 34(1), 25–42.

32. Jack, B. M., & Lin, H. S. (2017). Making learning interesting and its application to the science classroom. Studies in Science Education, 53(2), 137–164.

33. Jolliffe, I. T. (2002). Principal component analysis for special types of data (pp. 338-372). Springer New York.

34. Osborne, J., Simon, S., & Collins, S. (2003). Attitudes towards science: A review of the literature and its implications. International Journal of Science Education, 25(9), 1049–1079.

35. Owston, R., York, D. N., & Malhotra, T. (2019). Blended learning in large enrolment courses: Student perceptions across four different instructional models. Australasian Journal of Educational Technology, 35(5), 29–45.

36. Potvin, P., & Hasni, A. (2014). Analysis of the decline in interest towards school science and technology from grades 5 through 11. Journal of Science Education and Technology, 23, 784–802.

37. Raudenbush, S. W., & Bryk, A. S. (2002). Hierarchical linear models: Applications and data analysis methods

38. Renninger, K. A., Hidi, S., Krapp, A., & Renninger, A. (2014). The role of interest in learning and development.

39. Sample McMeeking, L. B., Weinberg, A. E., Boyd, K. J., & Balgopal, M.M. (2016). Student perceptions of interest, learning, and engagement from an informal traveling science museum. School Science and Mathematics, 116(5), 253–264.

40. Sinatra, G. M., Heddy, B. C., & Lombardi, D. (2015). The challenges of defining and measuring student engagement in science. Educational Psychologist, 50(1), 1–13.

41. Singh, K., Granville, M., & Dika, S. (2002). Mathematics and science achievement: Effects of motivation, interest, and academic engagement. The Journal of Educational Research, 95(6), 323–332.

42. Schmidt, J. A., Rosenberg, J. M., & Beymer, P. N. (2018). A person in context approach to student engagement in science: Examining learning activities and choice. Journal of Research in Science Teaching, 55(1), 19–43.

43. Taber, K. S. (2018). The use of Cronbach’s alpha when developing and reporting research instruments in science education. Research in Science Education, 48, 1273–1296.

44. Taylor, L. & Parsons, J. (2011). Improving student engagement. Current issues in education, 14(1).

45. Tröbst, S., Kleickmann, T., Lange-Schubert, K., Rothkopf, A., & Möller, K. (2016). Instruction and students’ declining interest in science: An analysis of German fourth-and sixth-grade classrooms. American Educational Research Journal, 53(1), 162–193.

46. Vedder Weiss, D., & Fortus, D. (2012). Adolescents’ declining motivation to learn science: A follow up study. Journal of Research in Science Teaching, 49(9), 1057–1095.

47. Waller, B. M., Peirce, K., Mitchell, H., & Micheletta, J. (2012). Evidence of public engagement with science: visitor learning at a zoo-housed primate research centre.

48. Wang, M. T., & Eccles, J. S. (2012). Adolescent behavioral, emotional, and cognitive engagement trajectories in school and their differential relations to educational success. Journal of Research on Adolescence, 22(1), 31–39.

49. Wang, M. T., & Holcombe, R. (2010). Adolescents’ perceptions of school environment, engagement, and academic achievement in middle school. American Educational Research Journal, 47(3), 633–662.

50. Wolf, S. J., & Fraser, B. J. (2008). Learning environment, attitudes and achievement among middle-school science students using inquiry-based laboratory activities. Research in Science Education, 38, 321–341.

